# KRAS Inhibition Reverses Chemotherapy Resistance Promoted by Therapy-Induced Senescence-like in Pancreatic Ductal Adenocarcinoma

**DOI:** 10.1101/2025.04.13.648164

**Authors:** Analia Meilerman Abuelafia, Patricia Santofimia-Castaño, Matias Estaras, Daniel Grasso, Eduardo Chuluyan, Gwen Lomberk, Raul Urrutia, Nelson Dusetti, Nicolas Fraunhoffer, Juan Iovanna

## Abstract

**Background:** Emerging evidence suggests that chemotherapy can accumulate senescent-like cells within tumor tissues, a phenomenon linked to therapy resistance. The aim of this study is to analyze the senescence-like state of after-treatment persistent cells associated with KRAS mutational status to offer a therapeutic strategy to target these cells in pancreatic ductal adenocarcinoma (PDAC).

**Experimental Design:** Three commercial cell lines and five patient-derived primary cell cultures with different KRAS statuses were studied following gemcitabine treatment. Senescence-like status was assessed using SA-β-gal, together with cell cycle regulators such as p21. Additionally, KRAS mutations were modulated using MRTX1133 and AMG-510, and the signaling pathways ERK and AKT were analyzed and modulated in vitro. Finally, p21 expression, associated with the senescence-like state, on patient outcomes and treatment response was analyzed in publicly available bulk RNA-seq and single-nucleus datasets.

**Results:** We observed an overexpression of p21 alongside an increase in SA-β-gal signal in response to gemcitabine treatment, indicating the induction of a senescence-like state. Specific inhibition of KRAS G12D or G12C mutations reduced SA-β-gal signal and sensitized PDAC cells to gemcitabine. Moreover, ERK inhibition but not AKT inhibition decreased SA-β-gal signal. Additionally, we characterized p21 expression levels in relation to patient outcomes and found that they are modulated by treatment.

**Conclusions:** This dual-targeted therapeutic strategy holds promises for overcoming the challenges posed by KRAS-driven cancers, particularly in addressing the formidable obstacle of pancreatic cancer.

**Highlights:**

➔ The accumulation of senescent-like cells in tumor tissues because of chemotherapy is linked to treatment resistance, specifically in pancreatic ductal adenocarcinoma (PDAC).
➔ Therapy-induced senescence-like in PDAC is influenced by the mutational status of KRAS, affecting the response to chemotherapeutic agents.
➔ The expression of the p21 is activated in response to therapy-induced senescence-like, particularly after gemcitabine treatment and is associated with patients outcome.
➔ Specifically inhibiting the mutated form of KRAS with small compounds can sensitize PDAC cells to gemcitabine and reduce SA-β-gal signal, acting through the ERK pathway but not the AKT pathway.
➔ The study presents a dual-targeted therapeutic strategy that combines KRAS inhibition with cytotoxic agents, potentially improving treatment efficacy and overcoming resistance in KRAS-driven cancers, such as PDAC.

## Introduction

Pancreatic ductal adenocarcinoma (PDAC) is one of the most lethal forms of cancer and it is projected to become the second cause of cancer-related death by 2030 (1). Despite advances in diagnosis and treatment in the past decade, the 5-year survival rate of PDAC is approximately 12%. The high mortality rate in PDAC underscores the shortcomings of conventional therapies, emphasizing the urgency for new therapeutic approaches. Surgical removal is viable in only a minority of cases due to the predominance of unresectable or metastatic tumors (2), with frequent relapse even in localized instances. Chemotherapy, particularly gemcitabine, either alone or in conjunction with nab-paclitaxel or mFOLFIRINOX regimen, stands as the primary treatment option following relapse or in cases of advanced disease. However, despite its widespread use, its effectiveness is hampered by the development of resistance (2–7).

Thus, in pancreatic cancer, resistance and not initial therapy represent the largest unmet need. In fact, PDAC cells manifest resistance to therapeutic drugs through a complex array of mechanisms, including the inactivation of drugs, alterations in drug metabolism, epigenetic changes, and modifications in drug targets (8). Preclinical studies suggests that chemotherapy can lead to the accumulation of senescent cells in tumor tissues (9). These senescent cells have been implicated in therapy resistance. Despite the ability of malignant tumors to evade senescence, they can still be forced to enter a senescent state using therapeutics leading to therapy-induced senescence (10,11). Conventional anticancer therapeutics, such as chemotherapy or radio-therapy, are known to induce senescence in cancer. On one hand, it is proposed that senescence can contribute to antitumor effects and treatment outcomes (12). Contrary, chronic accumulation of senescent cells can stimulate relapse and metastasis (13). For cancer cells carrying genomic alterations in cell cycle regulator genes, such as TP53 or CDKN2A, the cell cycle arrest associated with senescence is often unstable, allowing these cells to potentially resume proliferation. In this context, studies on colon cancer cells treated with cisplatin have demonstrated the expression of senescence markers (SA-β-gal and p21) while still maintaining DNA synthesis. These cells displayed resistance to chemotherapy, constituting treatment-persistent cells (14). This hybrid state is therefore referred to as a senescence-like state (15).

This study reports that PDAC therapy-induced senescence-like is mediated by KRAS mutational status, with KRAS mutated being more prone to develop this state which may limit the effectiveness of the chemotherapeutic agents that disrupt the DNA replication although not exclusively. Moreover, we demonstrate that mutated KRAS specific inhibition reverses resistance and SA-β-gal activity through the ERK-dependent pathway. This new knowledge bears relevance to clinical investigations that can benefit from more evidence-based experimental therapies to be further developed for the treatment of this dismal disease.

## Results

### Chemotherapeutic drugs promote a senescence-like state in a KRAS mutation-dependent manner in PDAC-derived cells

To evaluate the association between KRAS mutational status and the senescence-like sate as mediator of resistance in PDAC, we first determined the capability of the chemotherapeutic drugs, gemcitabine, 5-fluorouracil (5FU), oxaliplatin, SN38 (the active metabolite of irinotecan), and paclitaxel at 0.1 and 1 µM to induce therapy-induced senescence-like in three commercial pancreatic cancer cell lines (CCL) with different *KRAS* status. The analysis was focused on the after-treatment persisting cells, which displayed the highest treatment-associated tolerance. For this purpose, we developed a testing set of cell lines, which included MiaPaCa-2 (G12C) Panc-1 (G12D), and BxPC-3 (WT) cells. Subsequently, we repeated the same experiments in a validation cohort of patient-derived pancreatic cancer cells. After 72 hours of treatment, the persisting cells were assessed for senescence-associated β-galactosidase (SA-β-gal) assay (14). We find that gemcitabine induces the highest increase in the percentage of SA-β-gal positive cells in the mutated *KRAS* cells compared with wildtype (Figure 1A). We find that gemcitabine induced in 89.49%, 56.25% and 13.14% of positive SA-β-gal staining for MiaPaCa-2, Panc-1 and BxPC-3 respectively. Treatment with 5FU, oxaliplatin, SN38 or paclitaxel also showed activation of therapy-induced senescence-like, although with less intensity as presented in Figure 1B. These observations were validated in an independent set of five patient-derived primary cell culture (PDC), four mutated (PDAC001T (G12D), PDAC024T (G12D), PDAC054T (G12C), and PDAC022T (G12R) and one wildtype (PDAC064T) treated with gemcitabine, where the *KRAS* mutated PDCs displayed the higher increase in SA-β-gal positive cells (79.05%, 76.60%, 58.55% and 43.57% respectively) compared to *KRAS* WT (13.88%), as showed in Supplementary Figure 1. The same pattern was detected with SA-β-gal luminescence assay, where the KRAS mutated cells, MiaPaCa-2 and Panc-1 displayed a higher signal than the wild-type BxPC-3 cell after the treatment with gemcitabine (Figure 1C). Additionally, gemcitabine-persistent cells exhibited a statistically significant increase in nuclear area (Figure 1D). Thus, combined the results from both the testing set and patient-derived pancreatic cancer cell lines demonstrate that after treatment persistent cells displays features associated with a senescence-like state in PDAC and suggest KRAS status as a mediator of this process.

**Figure 1.**
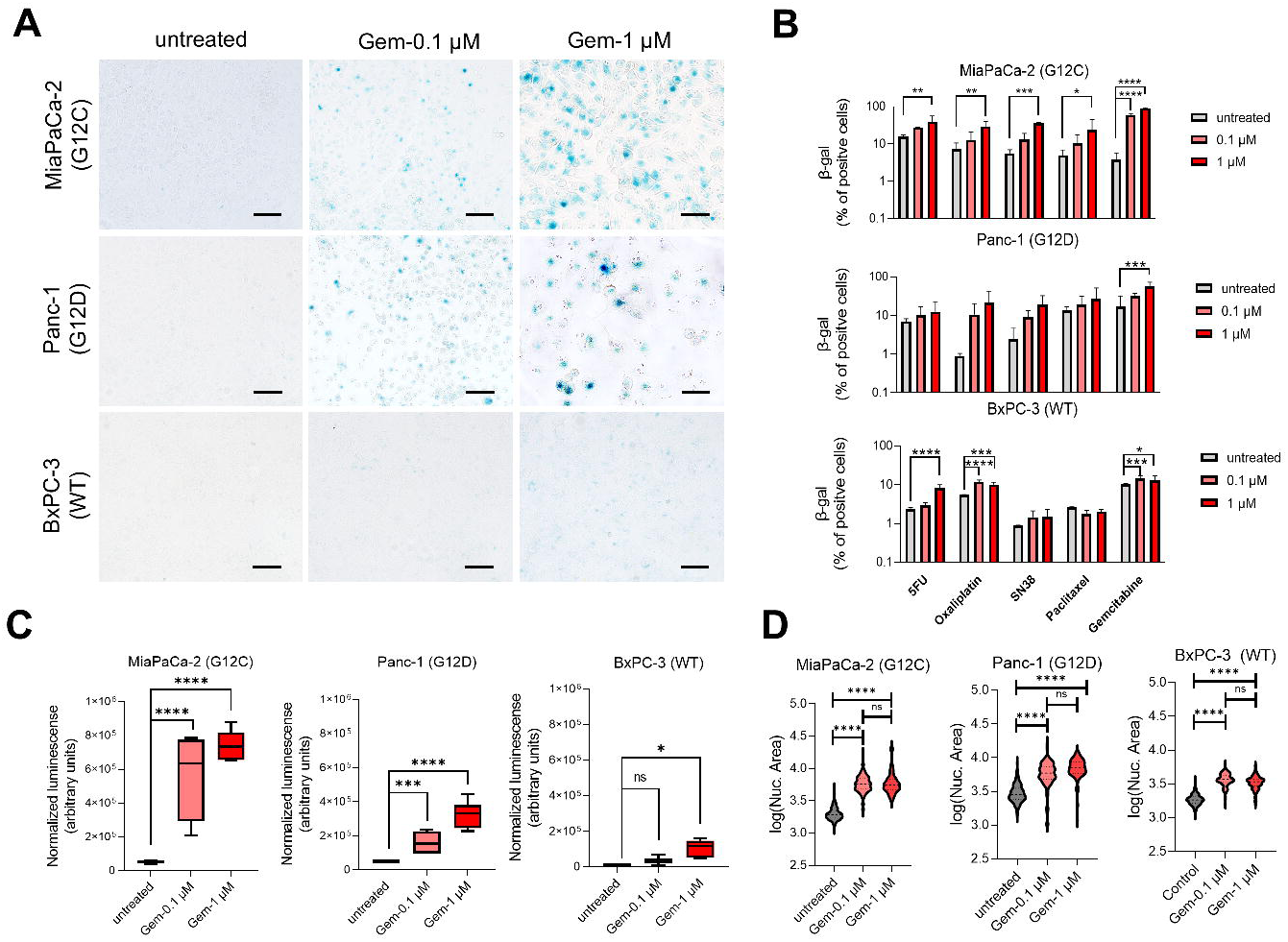
Increase of SA-β-galactosidase (SA-β-gal) levels after chemotherapeutic treatment in PDAC cells. A. Induction of SA-β-gal after the treatment with gemcitabine (72 hours) 0.1 µM and 1 μM. B. Percentage of SA-β-gal cell after the treatment with gemcitabine, 5-fluorouracil (5FU), oxaliplatin, SN38 (the active metabolite of irinotecan), and paclitaxel at 0.1 and 1 µM for 72 hours in MiaPaCa-2, Panc-1, and BxPC-3 cell lines. C. SA-β-gal expression measured by luminescence after the treatment by 72 hours with gemcitabine 0.1 µM and 1 μM. D. Measurement of nuclear area after the treatment by 72 hours with gemcitabine. Scale bars, 100 μm. \**P*<0.01, \*\**P*<0.01, \*\*\**P*<0.001, \*\*\*\**P*< 0.0001 (n = 3).

### PDAC senescence-like state is associated with p21 overexpression in treatment-persistent cells

Senescence-like state in cancer cells has been associated with a hybrid phenotype where the cells display markers of senescence but without complete cell cycle arrest (15). To further validate this resistant-associated phenotype, we assessed the expression levels of p21, p27, Ki67, CDK2, CDK4, and CDK6 in the persistent cell of the three CCL after 72 hours of treatment with gemcitabine at 0.1 and 1 µM). This experiment showed increased levels of the cell cycle inhibitor p21 in MiaPaCa-2 and Panc-1 cells whereas it decreases in BxPC-3 (see Figure 2A). Immunofluorescence microscopy using p21 antibodies validated these results as shown in Figure 2B and 2C. On the other hand, p27 decreases after the treatment in MiaPaCa-2 and Panc-1 cells while remaining unchanged in BxPC-3 (Figure 2A). Regarding the positive regulators of the cell cycle, we find an increase in Ki67 with the treatment in mutated KRAS. Similarly, we observe that CDK2 increases in mutated *KRAS* cells *vs.* wildtype. Moreover, the levels of CDK4 and CDK6 did not change (Figure 2A). In addition, we evaluated the expression of the epithelial-to-mesenchymal transition factor ZEB1, known to be activated in treatment-resistant cells, and found an increase after the cytotoxic treatment in mutated *KRAS* cells but not in wildtype which is almost undetected (Figure 2A). Collectively, these findings show that cytotoxic treatment induces a senescence-like state in a p21-dependent manner in the persistent cell and is associated to *KRAS* mutational status.

**Figure 2.**
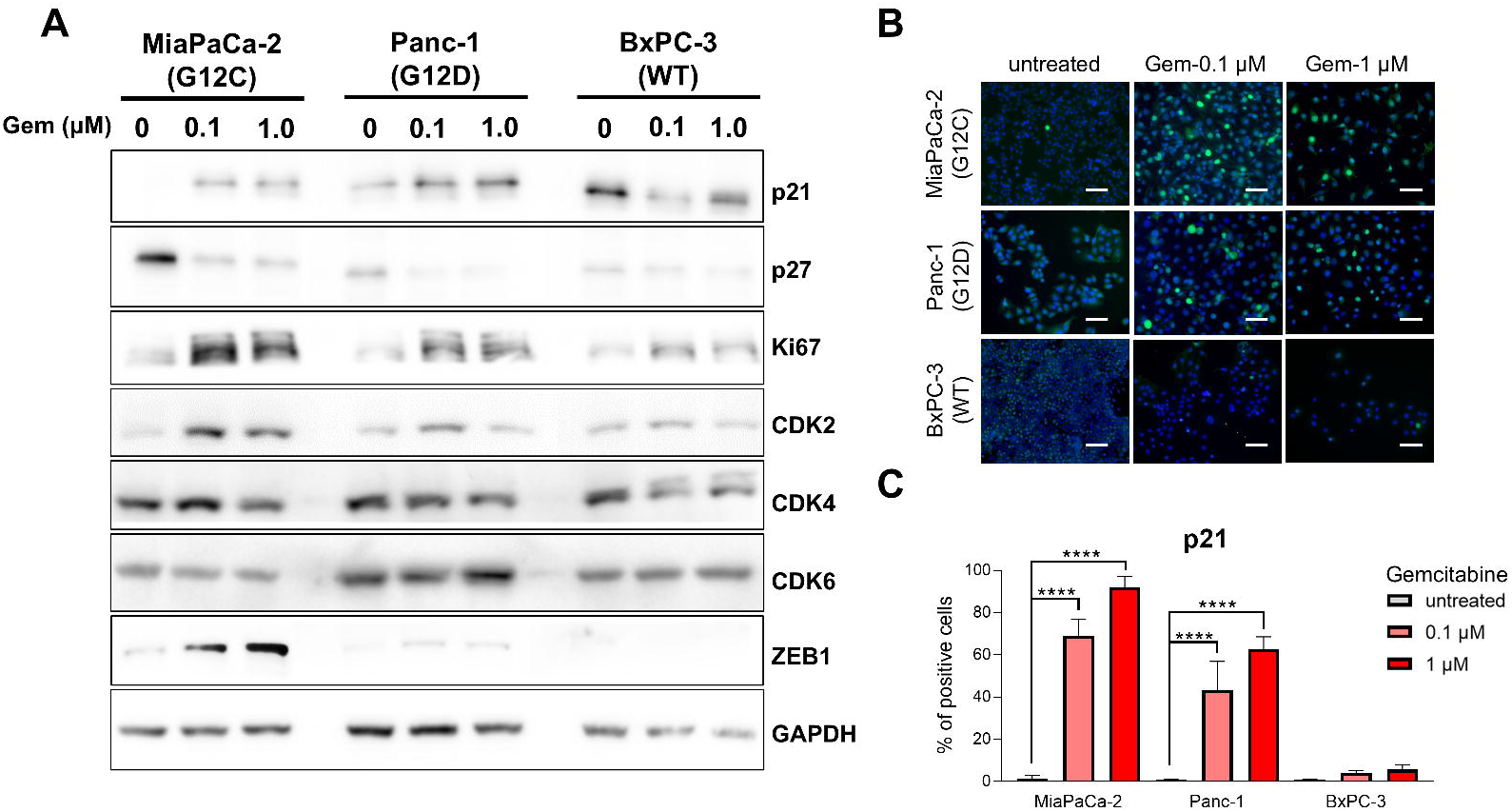
Senescence-like PDAC cells overexpress p21 after gemcitabine treatment. A. Immunoblotting of the cell cycle and epithelial-mesenchymal transition proteins after the treatment with gemcitabine at 0.1 µM and 1 μM for 72 hours in MiaPaCa-2, Panc-1, and BxPC-3 cell lines. B. Immunofluorescent staining of p21 in MiaPaCa-2, Panc-1, and BxPC-3 after treatment with gemcitabine at 0.1 µM and 1 μM. C. Percentage of the p21 stained cells. Scale bars, 200 μm. \*\*\*\**P*<0.0001 (n = 3).

### KRAS inhibitors antagonize therapy-induced senescence-like and sensitize persistent cells to gemcitabine

Next, to further confirm the dependence between *KRAS* mutational status and the senescence-like state induced by cytotoxic compounds, we inhibited KRAS activity in the CCL with two selective inhibitors, MRTX1133 and AMG-510 at 1 μM, designed to target G12D and G12C mutations, respectively. We find that ERK phosphorylation induced by cytotoxic treatment was abolished by the KRAS inhibitors in MiaPaCa-2 and Panc-1 but not in BxPC-3 (Figure 3A, 3B and 3C). We investigated whether KRAS inhibitors could influence therapy-induced senescence-like. To address this question, we treated MiaPaCa-2, Panc-1, and BxPC-3 cells with gemcitabine at 0.1 μM in combination with either KRAS inhibitors AMG-510 or MRTX1133 for MiaPaCa-2 and Panc-1, respectively. Subsequently, we measured SA-β-gal activity and found that AMG-510 or MRTX1133 alone did not induces senescence-like in these cells (Figure 3D, 3E and 3F). SA-β-gal activation induced by gemcitabine in *KRAS*-mutated cells was completely reversed by AMG-510 or MRTX1133 (Figure 3G and 3H). Moreover, treatment of BxPC-3 with gemcitabine alone or together with MRTX1133 does not modify the SA-β-gal signal (Figure 3I).

**Figure 3.**
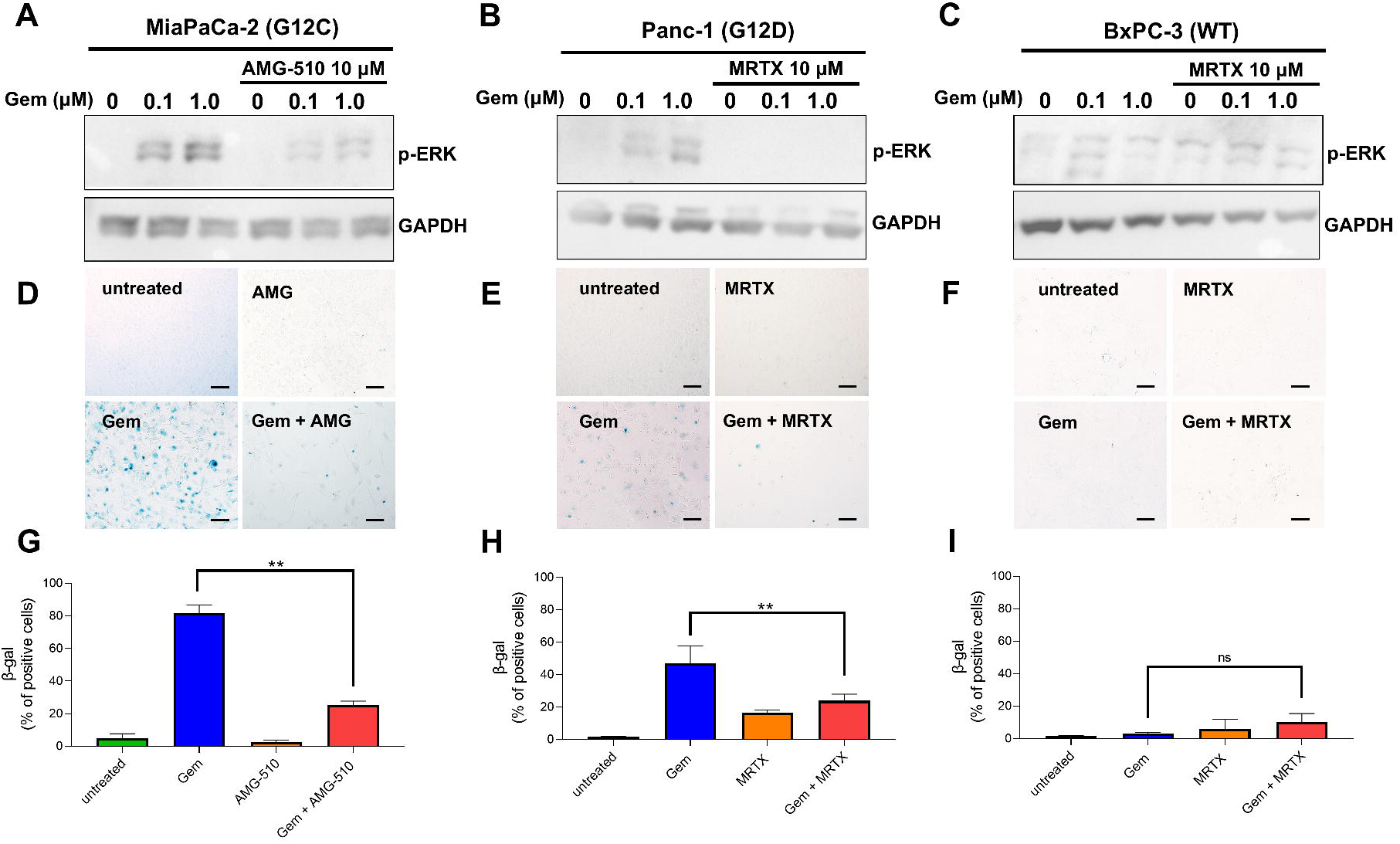
Reduction of SA-β-gal levels after mutated KRAS inhibition. KRAS inhibitors AMG-510 and MRTX1133 reduce the phosphorylation of ERK in MiaPaCa-2 and Panc-1 (A and B) but not in BxPC-3 (C). SA-β-gal staining levels after the treatment with gemcitabine in MiaPaCa-2 (D) and Panc-1 (E) without significant effect on BxPC-3 (F). G, H and I. Percentage of the SA-β-gal stained cells. Scale bars, 100 μm. \*\**P*<0.01, (n = 3).

Finally, we explored the impact of silencing therapy-induced senescence-like on gemcitabine resistance. Consequently, we exposed Panc-1 or MiaPaCa-2 cells to escalating concentrations of gemcitabine in conjunction with 1 µM of MRTX1133 or AMG-510, respectively, and assessed their cell viability. Additionally, we treated two primary cells cultures of PDAC carrying *KRAS* mutations (PDAC001T and PDAC024T) and one wildtype (PDAC064T) with gemcitabine alone or in combination with KRAS inhibitors. Moreover, we treated PDAC22T PDC, carrying a G12R mutation, with the combination of gemcitabine and MRTX1133 as a negative control, knowing that MRTX1133 does not target this mutation. The CCL and PDC carrying KRAS with mutations, G12C or G12D, showed a significant and systematic decrease in their area under de curve (AUC) compared to the wildtype and G12R mutated cells, in presence of the inhibitors (32%±10% *vs.* 11%±1%, *P*=0.026) as presented in Figure 4. It is important to note that neither AMG-510 nor MRTX1133 exhibited cytotoxic activity at the concentration used (Supplementary Figure 2). The synergy between KRAS inhibitors and gemcitabine was further confirmed with time-lapse analysis for 5 days, where the combination of AMG-510 or MRTX1133 with gemcitabine resulted in enhanced cytotoxicity, particularly in cells harboring *KRAS* G12D or G12C mutations, compared with the *KRAS* WT or G12R mutated cells (Figure 5). All together, these results demonstrate the positive association of KRAS mutational status and the response to gemcitabine in treatment-persistent PDAC cells.

**Figure 4.**
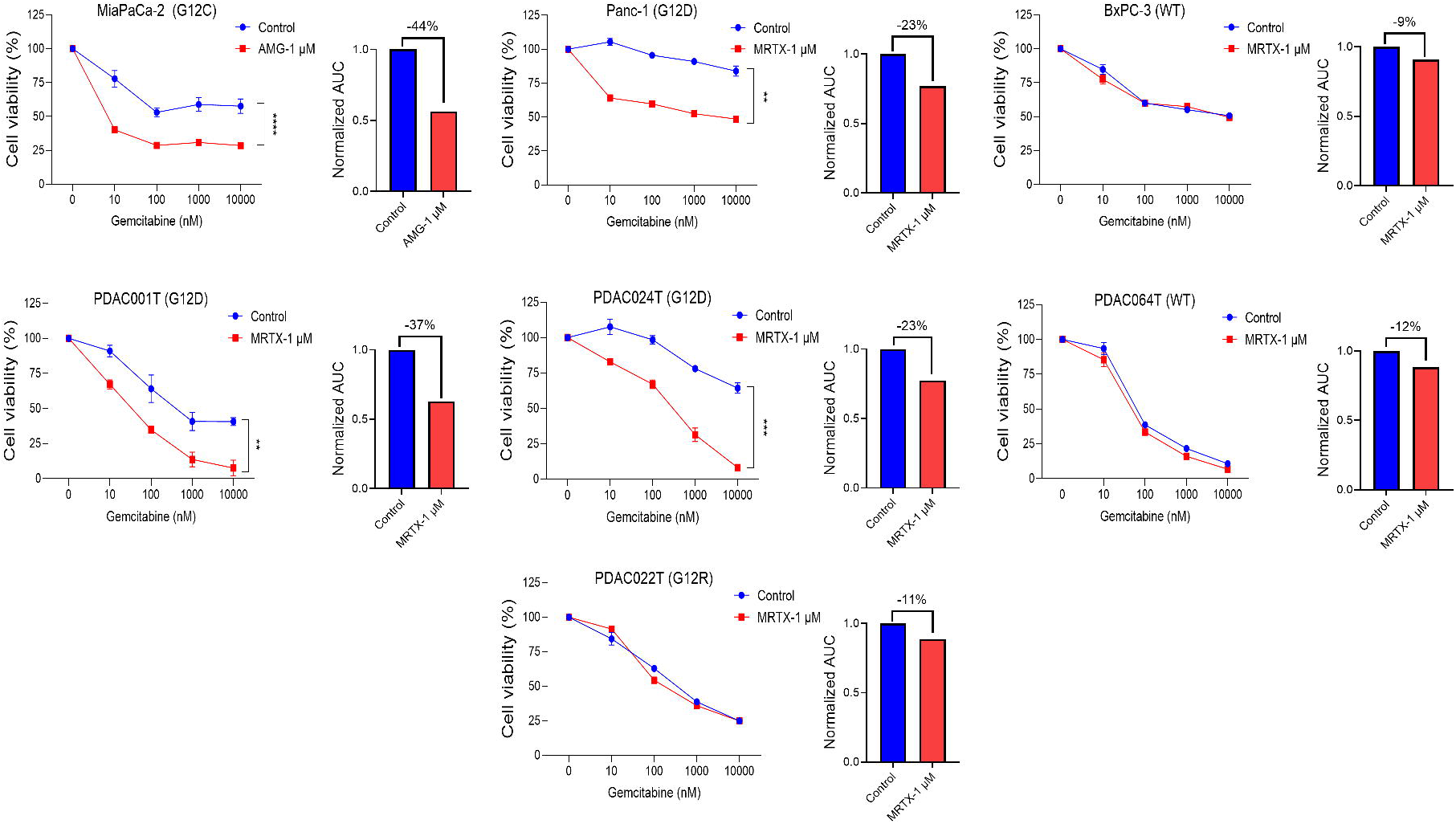
Inhibition of mutated KRAS sensitize PDAC cells to gemcitabine treatment. Three commercial cell lines MiaPaCa-2, Panc-1, and BxPC-3 and four patient-derived primary cell cultures were treated with increased concentrations of gemcitabine (from 0.01 μM to 10 μM) in combination with 1 μM of KRAS inhibitors, MRTX1133 (directed against G12D) or AMG-510 (directed against G12C) for 72 hours. Cell viability was measured with PrestoBlue. Each experiment was repeated at least three times. Values were normalized and expressed as the percentage of the control (vehicle). Additionally, the reduction in area under the curve (AUC) is shown. \*\**P*< 0.01, \*\*\*\**P*< 0.0001.

**Figure 5.**
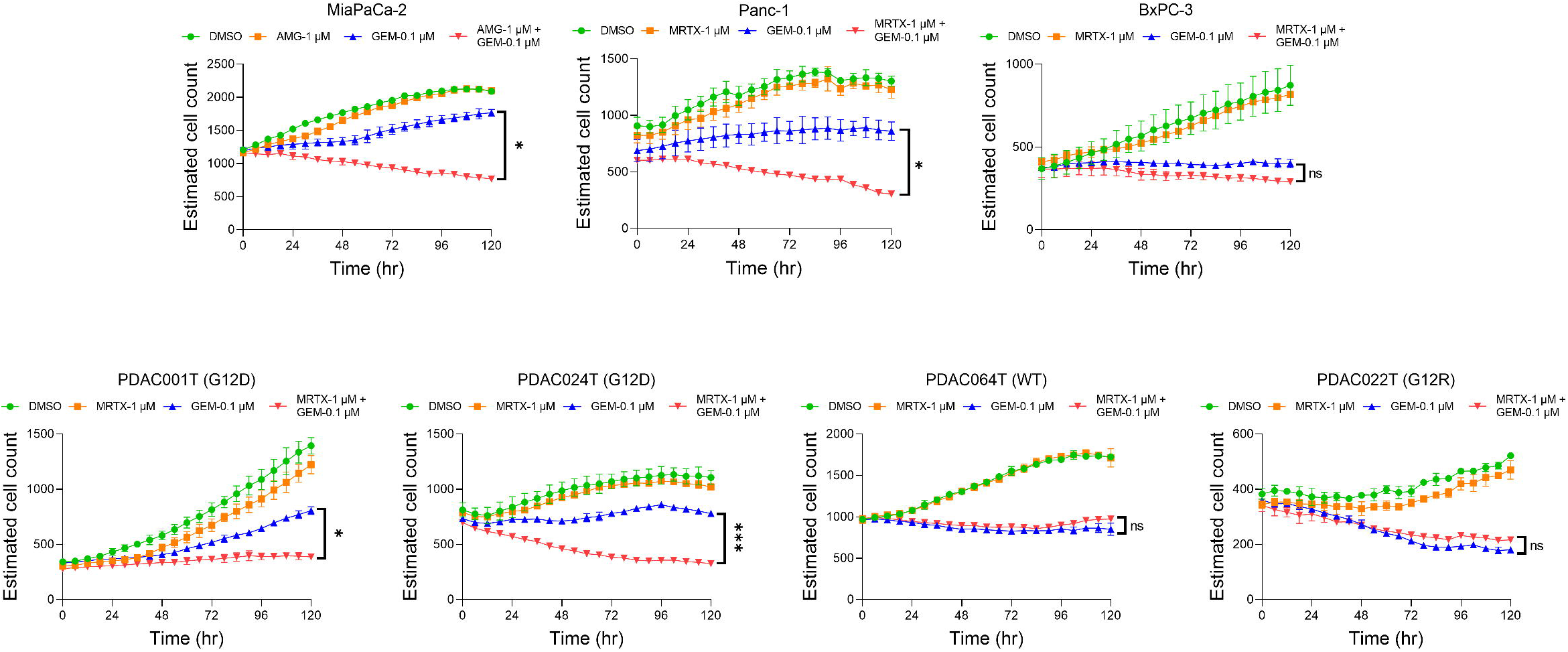
Time-lapse experiment analyzing the sensitize effect KRAS inhibition on gemcitabine treatment. Incucyte video microscopy measured the cell viability of 3 commercial cell lines MiaPaCa-2, Panc-1, and BxPC-3 and four patient-derived primary cell cultures, treated with gemcitabine 0.1 μM, 1 μM of KRAS inhibitors, MRTX1133 (directed against G12D) or AMG-510 (directed against G12C) or the combination for 120 hours. A control with DMSO (vehicle) was maintained. \**P*< 0.05, \*\*\**P*< 0.001, ns, not significant (n=2).

### Therapy-induced senescence-like is dependent of ERK-signaling pathway

Next, we explored the impact of gemcitabine treatment on the activation of ERK and AKT pathways. As illustrated in Figure 6A, phosphorylation of ERK (p-ERK) was markedly increased in MiaPaCa-2 and Panc-1 cells following gemcitabine treatment, while no significant change was observed in BxPC-3 cells. Notably, the phosphorylation of AKT at activation sites S473 and, especially, on T308 was elevated in *KRAS*-mutated MiaPaCa-2 and Panc-1 cells, but not in wildtype BxPC-3 cells. In addition, phosphorylation of PTEN, the phosphatase regulating AKT activation, at amino acid S380 exhibited a notable decrease in *KRAS*-mutated cells compared to wildtype cells. Moreover, phosphorylation of PDK1 at amino acid S241 reveals a strong inhibition of its activity. Lastly, while wildtype BxPC-3 cells show no changes in p-AKT (S473), p-AKT (T308), and p-PTEN (S380), total AKT and p-PDK (S241) accumulate after the treatment. Lastly, immunofluorescence microscopy also shows the activation of p-ERK in response to gemcitabine (Figure 6B and 6C). Next, since both ERK and AKT signaling pathways are activated by therapy-induced senescence-like, we investigated whether ERK, AKT or both mediated this process. We treated cells with gemcitabine alone or together with increasing concentrations of ERK (SCH772984) or AKT (ipatasertib) inhibitors and measured the SA-β-gal signal. We observed that SA-β-gal signal decreases in MiaPaCa-2 and Panc-1 cells treated with SCH772984 while no changes were observed in BxPC-3 cells (Figure 7A). On the contrary, treatment of the cells with ipatasertib reveals the lack of inhibition of SA-β-gal signal (Figure 7B). Altogether, these results suggest that the therapy-induced senescence-like is mediated by ERK but not by AKT pathways.

**Figure 6.**
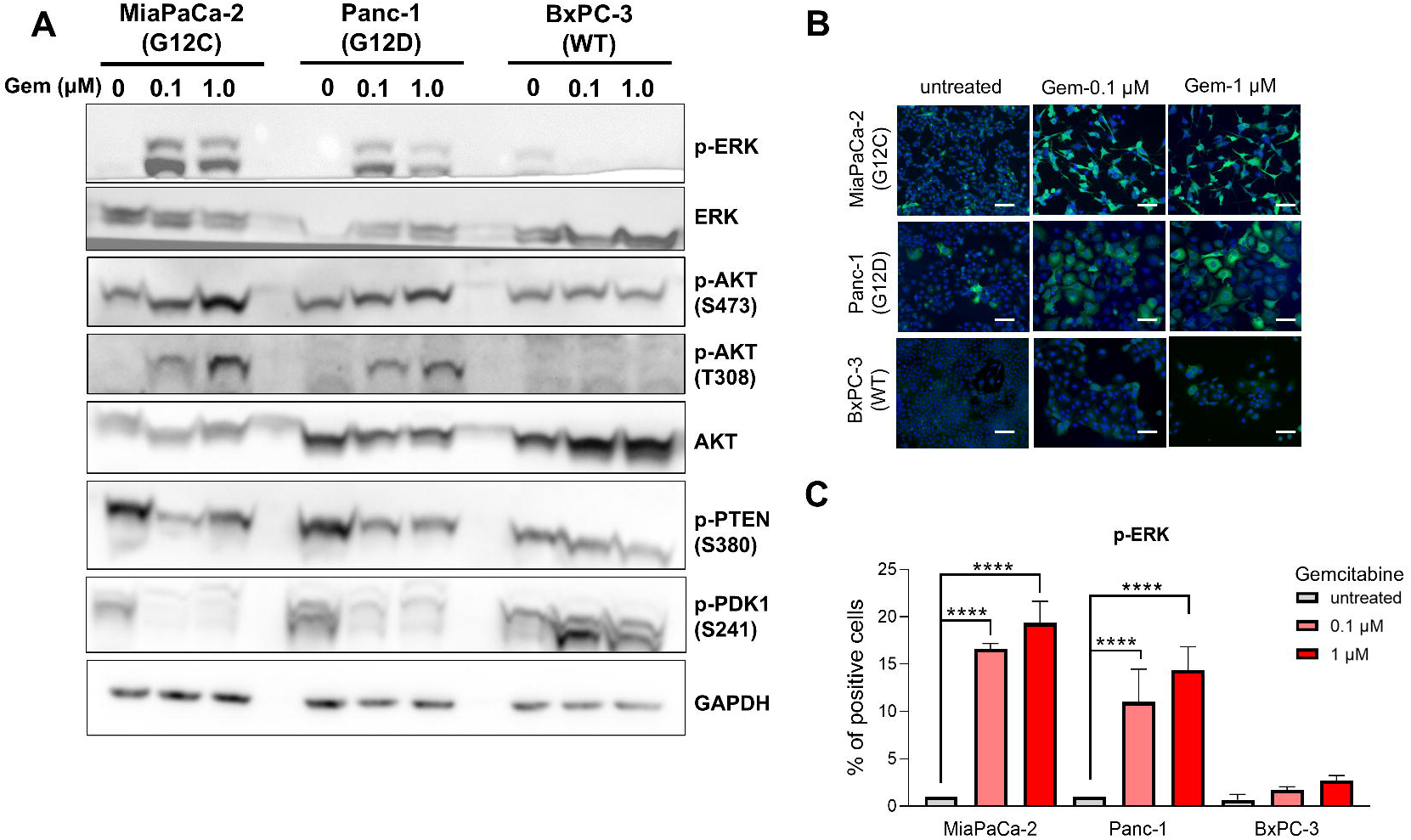
Characterization of KRAS pathway in PDAC cells after the treatment with gemcitabine. A. Immunoblotting of ERK and AKT pathway in MiaPaCa-2, Panc-1, and BxPC-3 after treatment for 72 hours with gemcitabine at 0.1 µM and 1 μM. B and C. Immunofluorescent staining of p-ERK and quantification of positive cells for p-ERK labeling after the treatment with gemcitabine at 0.1 µM and 1 μM for 72 hours in MiaPaCa-2, Panc-1, and BxPC-3. Scale bars, 200 μm. \*\*\*\**P*<0.0001, (n = 3).

**Figure 7.**
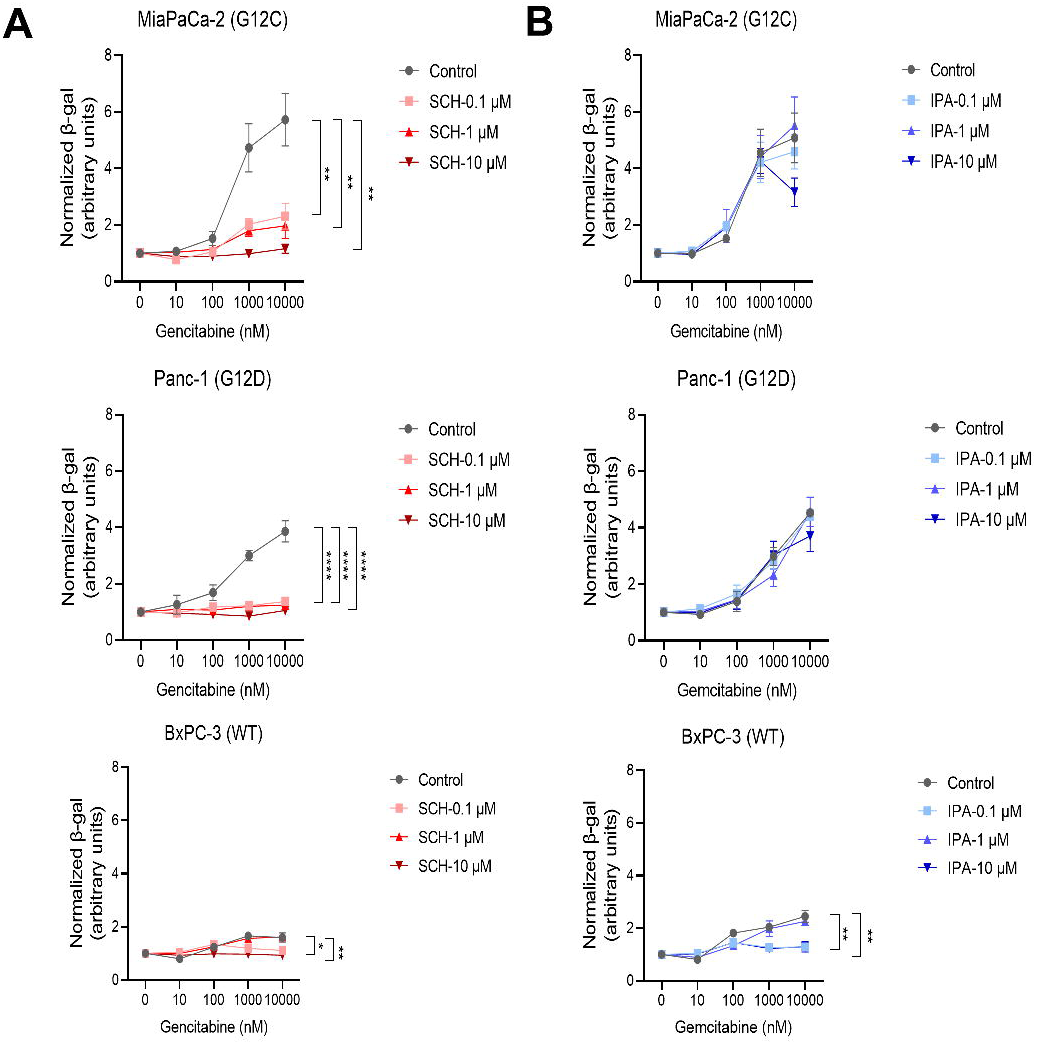
ERK inhibitor reduces the levels of SA-β-gal in presence of gemcitabine. SA-β-gal level was measured by luminescence assay after the treatment with gemcitabine in increasing concentrations (from 0.01 μM to 10 μM) in combination with (A) the ERK (SCH772984; SCH) and (B) AKT (Ipatasertib; IPA) inhibitors (0.1 μM, 1 µM and 10 µM). \**P*<0.05, \*\**P*<0.01, \*\*\*\**P*<0.0001. (n = 3).

An important point to highlight is that, like gemcitabine treatment, the effect of 5-FU, oxaliplatin, irinotecan, and paclitaxel on p-ERK activation has been previously documented in other tumors-derived cells (16–19). We also demonstrated that SA-β-gal increase of signal depends on p-ERK activation, as blocking p-ERK with KRAS inhibitors prevents its detection, as shown in Figure 3. The induction of SA-β-gal by cytotoxic agents has also been previously reported in other tumors-derived cells (20,21). Mechanistically, the activation of p-ERK following treatment with cytotoxic drugs occurs as a response to genotoxic stress and DNA damage. Additionally, p-ERK stimulates the production of SASP (senescence-associated secretory phenotype), which reinforces autocrine senescence and alters the tumor microenvironment, leading to inflammation and tumor resistance (22).

### Characterization of senescence-like cells associated with p21 expression in patients with PDAC

As previously mentioned, the senescence-like phenotype observed in treatment-persistent cell populations is dependent on p21 expression. To further investigate this, we assessed the association of p21 with prognosis in PDAC patients using bulk RNA-seq data, as well as its dynamics before and after treatment at the single-nucleus level. Initially, the TCGA PAAD and Puleo (23) cohorts were analyzed to evaluate the impact of p21 expression on overall survival (OS). Patients with high p21 expression had significantly shorter OS compared to those with low p21 expression in both the TCGA PAAD (12.7 vs. 21.1 months, HR: 0.53; 95% IC: 0.31– 0.89, *P*=0.020, Figure 8A) and Puleo (18.5 vs. 30.4 months, HR: 0.52; 95% IC: 0.37–0.73, *P*<0.001, Figure 8B) cohorts. To further validate the association between p21 and the senescence-like phenotype, we performed single-sample enrichment analysis using the EpiSen signature from Gavish et al. (24). High p21 group exhibited a significantly higher EpiSen score compared to the low p21 group in both the TCGA PAAD (Figure 8A) and Puleo (Figure 8B) cohorts. Lastly, we assessed the effect of treatment on p21 expression at the single-nucleus level using the Hwang et al. (25) cohort. The analysis focused on previously annotated malignant cells, comprising a total of 47,832 cells. Consistent with our previous observations in CCL and PDC models, treated cells exhibited higher p21 expression and an increased EpiSen score compared to untreated cells (Figure 8C). These results validate the association between p21 and the senescence-like phenotype in PDAC patients, highlighting its prognostic significance and the role of treatment as an inducer of this cellular state.

**Figure 8.**
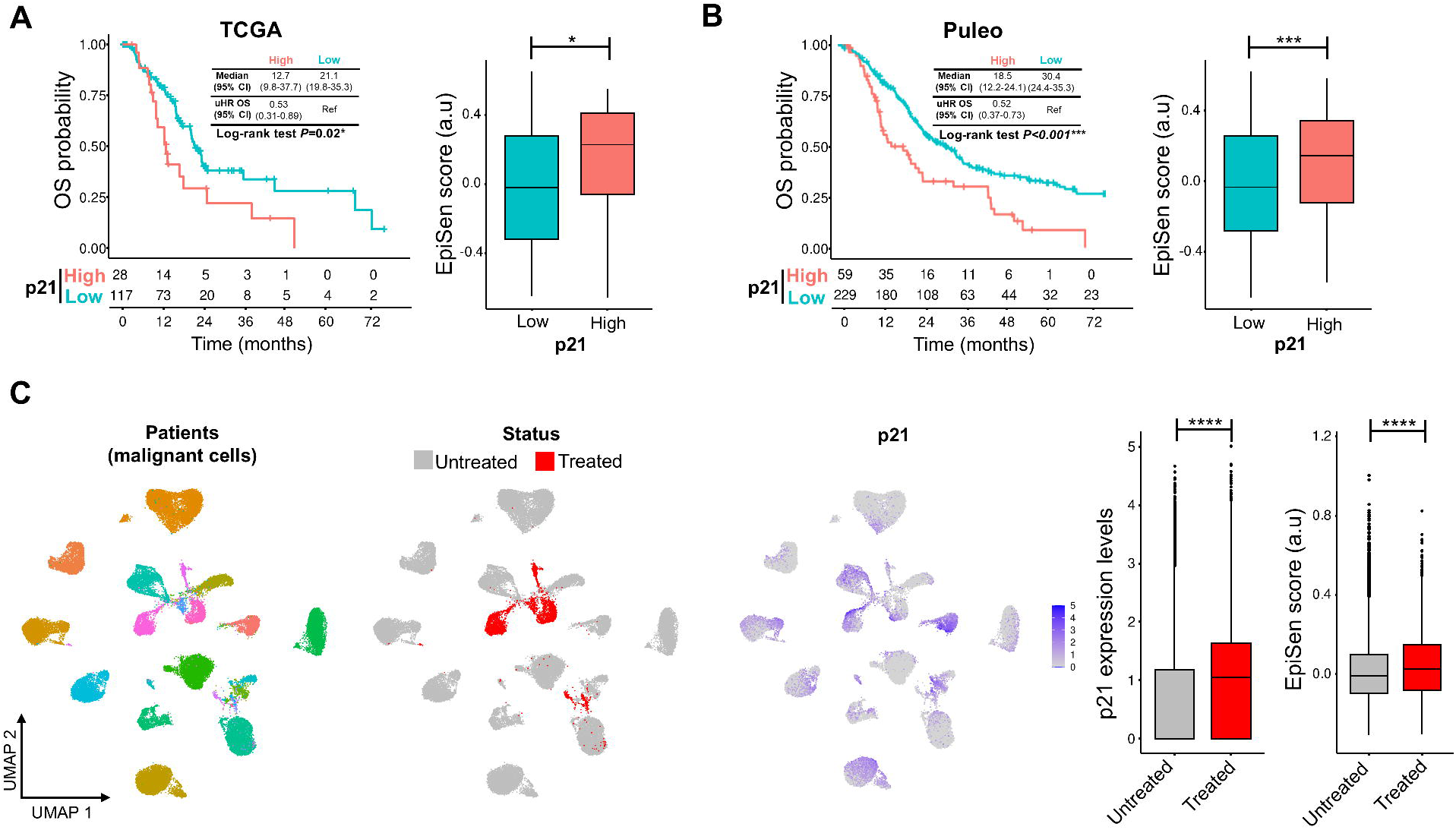
Expression levels of p21 as mediator of the senescence-like state is associated with patient’s outcome and persistent cells after treatment. A. Survival analysis of p21 levels associated with overall survival (OS) and senescence-like signature EpiSen in TCGA-PAAD (A) and Puleo (B) cohorts. C. Expression of p21 and score of EpiSen signature before and after treatment in single nucleus Hwang cohort. \**P*<0.05, \*\*\**P*<0.001, \*\*\*\**P*<0.0001. HR, Hazard ratio, IC, interval of confidence.

## Discussion

The current manuscript contributes with new knowledge applicable to current clinical efforts for treating pancreatic cancer, through a combination of targeted KRAS inhibition and chemotherapy. Several mechanisms contribute to the resistance of PDAC to cytotoxic treatments, among which one of the most significant is therapy-induced senescence-like (26). Mutated *KRAS* plays a pivotal role in both development and progression of PDAC (27), as well as in conferring resistance to treatment through the activation of therapy-induced senescence-like as demonstrated in this work. Senescence-like entails a state of activation of cell cycle regulators such as p21 but without a complete suppression of DNA synthesis activity, rendering senescent-like cells insensitive to some cytotoxic drugs. In this paper we describe that therapy-induced senescence-like is prone to be activated in PDAC cells carrying *KRAS* mutations, but not on those with a wildtype sequence, and that treatment with specific KRAS inhibitors hinders senescence-like and switching the cells toward a more sensitive phenotype. Expression of the cell cycle inhibitor p21 is activated in cells carrying *KRAS* mutations in response to cytotoxic agents to activate the therapy-induced senescence-like. Remarkably, ERK-but not AKT-dependent signaling is mediating the therapy-induced senescence-like.

Several studies have demonstrated that KRAS inhibition can sensitize PDAC cells to gemcitabine treatment both in vitro (28,29) and in vivo (30–32) Additionally, the RAF/MEK/ERK pathway has been implicated in resistance to Gemcitabine, and its inhibition has been shown to enhance the drug’s efficacy (33–35). Given this evidence, the hypothesis that KRAS inhibition enhances Gemcitabine sensitivity via suppression of the RAF/MEK/ERK pathway is not only plausible but likely a critical mechanism underlying treatment resistance. A deeper understanding of these resistance pathways is essential for developing more effective therapeutic strategies, potentially transforming the treatment landscape for PDAC.

The synergy between drugs is highly dependent on the models used and the specific setup of the study protocol. The study by Gulay et al. (36) and the study conducted by Dr. Schneider’s group (37). report no synergic effect between the KRAS inhibitor MRTX1133 and gemcitabine. It is important to note that even when using the same cell line, results can vary due to differences in culture conditions, cellular heterogeneity, and biological adaptability. Factors such as media composition, cell density, passage number, and experimental methodology can influence cellular responses. Additionally, genetic and epigenetic variability within the cell population, along with the activation of adaptive mechanisms, can alter treatment sensitivity. These differences highlight the importance of standardizing protocols and conducting replications to ensure experimental reproducibility. Specifically, no synergistic effect was identified between gemcitabine and Sotorasib, a KRAS^G12C^ inhibitor, when using the MiaPaCa-2 cell line. This raises the possibility that the synergy may depend on specific components of the inhibitor’s structural framework or molecular interactions that are not apparent in this context. Furthermore, in the clinical trial published by Infante and collaborators (38), in which they applied the trametinib, an oral MEK inhibitor, in combination with gemcitabine for patients with PDAC, they didn’t found any synergism. In contrast, in the Hoffer’s report (39), a synergic effect with a MEK inhibitor was observed in several but not in all PDAC-derived cells. Altogether, these findings suggest that achieving synergy between gemcitabine and mutated KRAS forms likely involves intricate molecular mechanisms or factors that currently fall outside the bounds of our rational understanding.

While the data obtained from comparing KRASwt and KRASmut cell lines are statistically significant, we acknowledge that the limited number of models used in this study restricts the ability to generalize our conclusions to a broader range of biological contexts. Our results show a clear and reproducible difference between the responses observed in KRAS-mutant cells and those with wild-type KRAS, suggesting a relevant role of KRAS in regulating cellular response to treatment. However, the inclusion of only two KRASwt models compared to five KRASmut models implies that, although the observed trends are robust within our dataset, they cannot be considered definitive evidence of a universally applicable mechanism. To strengthen these conclusions, future studies should incorporate a larger number of KRASwt cell lines and, whenever possible, isogenic models, where the only differential variable is the presence or absence of the KRAS mutation. This approach would allow for a more precise evaluation of the impact of KRAS on treatment response modulation while ruling out potential effects associated with other genetic differences between the cell lines used.

KRAS, a predominant member of the RAS family, stands as the most frequently mutated oncogene in human PDAC, accounting for approximately 95% of cases. Mutations in *KRAS* instigate its constitutive activation, consequently triggering downstream signaling pathways like RAF-MEK-ERK and PI3K-AKT-mTOR. These pathways promote cell proliferation and equip cancer cells with capabilities to evade apoptosis. Historically considered “undruggable”, KRAS saw a breakthrough with the discovery of the first covalent inhibitor targeting the G12C mutation (40). However, G12C mutations, prevalent in non-small cell lung cancer, are relatively rare in PDAC. Instead, PDAC predominantly harbors other KRAS mutations such as G12D and G12V. Inhibitors targeting G12D mutation (e.g., MRTX1133) have been published in 2022 (33,41) and those addressing the pan-KRAS were developed just one year later revealing the pharmaceutical interest in this field. The pan-KRAS inhibitor BI-2493 and its structural analogue BI-2865, that was optimized for *in vivo* administration, effectively hindered nucleotide exchange, thereby preventing the activation of wild-type *KRAS* and a diverse array of *KRAS* mutants, encompassing G12A/C/D/F/V/S, G13C/D, V14I, L19F, Q22K, D33E, Q61H, K117N, and A146V/T. The specific observation of its impact on downstream signaling and proliferation was specifically observed in cancer cells harboring mutant *KRAS*. Remarkably, treatment with the drug significantly inhibited the growth of *KRAS* mutant tumors in mice, without eliciting any obvious adverse effects on animals (42). Nonetheless, the therapeutic efficacy of KRAS inhibitor monotherapy is hindered by the emergence of resistance mechanisms (43). While these compounds were originally developed with the aim of eradicating cancer cells, our proposal suggests utilizing this type of drug to impede senescence-like and enhance its synergy with cytotoxic agents in the treatment of resistant PDAC, giving a repositioning strategy.

In our analysis, we found that p21 is activated in response to gemcitabine treatment to play a main role in development of senescence-like (44). Surprisingly, alongside p21 activation, we observed concurrent activation of Ki67 and CDK2, which typically drive cell cycle progression, seemingly contradicting senescence induction. Notably, while some cell proliferation associated proteins such as Ki67 and CDK2 were activated, others, such as CDK4 and CDK6, remained unchanged, suggesting a non-canonical senescence pathway. The cell cycle is a complex process that requires successive involvement of multiple proteins, and only when all these components function properly can cellular proliferation occur. An increase in Ki-67 does not necessarily indicate an increase in proliferation. For instance, as demonstrated over 13 years ago (45), the co-overexpression of p21 and Ki-67 systematically occurs in the cells of head and neck squamous cell carcinoma tumors. Treatment with cytotoxic drugs induces a reduction in the number of viable cells. In this work, we also demonstrated that treatment with cytotoxic agents induce the phosphorylation of ERK1/2, and through this signaling pathway, the cell cycle could be apparently activated. Additionally, the induction of epithelial-mesenchymal transition (EMT) by ZEB1 was notably heightened during gemcitabine treatment (46). However, the precise alterations in the expression of key genes during therapy-induced senescence-like warrant further investigation. Notably, these changes in gene expression occur only in mutated *KRAS* cells.

Data from this work revealed some surprising results. Firstly, as referred above, we observed that in therapy-induced senescence-like there is an apparent opposite effect on the cell cycle through, from one hand, the over-expression of cell cycle inhibitor p21 and, on the other hand, the increasing of the promoting cell cycle factors CDK2 and Ki67. This can be interpreted as a response by the cell to have an equilibrium between growth arrest and promotion in response to the treatment with cytotoxic agents. The second fact is that in BxPC-3 cell, which is carrying a wildtype KRAS with an in-frame oncogenic (V600E) mutation on the BRAF, which may explain its transformed character, did not activate its downstream ERK1 and 2 factor in response to chemotherapy. The V600E point mutation promotes cell proliferation via the abnormal activation of the Ras/MAPK signaling pathway (47). However, despite the RAS/MAPK pathway being expected to be activated downstream of BRAF due to the V600E mutation, ERK remains inactive as presented in Figure 5. This could imply that ERK activation is dependent on a mutated form of KRAS through a non-canonical pathway for ERK activation in the case of therapy-induced senescence-like. Ramaker et al. (48) report the results of a CRISPR-based strategy in which authors investigated the gene pathways that were over-represented among those exhibiting drug resistance or sensitivity. Their analysis revealed several pathways associated with patient survival, including chromatin remodeling, hemidesmosome assembly, and without surprise the p-ERK signaling. These pathways were shown to promote drug resistance when activated and confer drug sensitivity when inhibited, underscoring their potential as therapeutic targets, particularly the p-ERK pathway. Finally, given that SA-β-gal staining is associated with increased lysosomal content (49), it is possible that the sensitization to gemcitabine observed after KRAS inhibition is not exclusively due to a reduction in senescence-like but rather to the blocking of autophagic and/or lysosomal functions. Hashimoto et al. (50), using drug concentrations comparable to those employed in our study, demonstrated that gemcitabine induces autophagic flux in both Panc-1 (KRAS^G12D^) and BxPC3 (KRAS^wt^) cells. This finding suggests that the SA-β-gal staining observed in our study may depend primarily on senescence-like rather than autophagy, as it is associated with treatment using KRAS inhibitors. This same phenomenon has also been documented with 5-fluorouracil (5-FU). The role of autophagy in cellular senescence-like is complex and remains a subject of intense scientific debate. On the one hand, autophagy has been linked to promoting senescence-like, particularly by facilitating the synthesis of the senescence-associated secretory phenotype (SASP) and enabling cells to bypass oncogene-induced senescence-like. On the other hand, autophagy acts as a cellular maintenance mechanism, clearing damaged macromolecules and organelles that might otherwise trigger senescence-like. This anti-senescence function of autophagy has been demonstrated in various contexts, including stem cells and cardiomyocytes (51–53). It is important to note that in vivo studies, particularly those employing transgenic mouse models, suggest that autophagy declines with aging. This decline is associated with the accumulation of damaged organelles, increased cellular senescence-like, and a higher cancer risk (54). These findings highlight the complexity of the relationship between autophagy and senescence-like and suggest that this interplay is dynamic, context-dependent, and multifaceted. The controversies surrounding this topic are widely discussed in the scientific community (55,56). Current debates underscore the need to unravel the specific molecular mechanisms and contextual roles of autophagy in establishing a senescent-like phenotype. In conclusion, the relationship between autophagy and senescence-like is highly intricate and remains far from fully understood. While this topic is beyond the scope of the present study, it represents a critical area for future research, with implications spanning aging, cancer biology, and the development of novel therapies.

In summary, our findings highlight a novel rational approach based on the treatment of PDAC cells with a cytotoxic agent while simultaneously impeding senescence-like through inhibiting mutant KRAS activity. Our results reveal a notable enhancement in cell death under these conditions underscoring its potentially impact on improving the efficacy of chemotherapy agents. In this manner, by combining new targeted therapies using KRAS inhibitors with conventional anticancer agents, we may harness effects that amplify the cytotoxicity against PDAC cells. Thus, this dual-targeted therapeutic approach holds promise for addressing the challenges posed by KRAS-driven cancers, offering a nuanced and effective strategy to combat pancreatic cancer.

## Limitations of the Study

Despite the significant findings presented in this study regarding KRAS inhibition and its impact on chemotherapy sensitivity in PDAC, several limitations must be considered when interpreting the results.

Limited Number of Cell Models: One of the main challenges of this study is the restricted number of cell models used. The study included five KRAS-mutant (KRAS^mut^) models and only two wildtype KRAS (KRAS^wt^) models. Although the data obtained show significant differences between the two groups, the disproportionate number of models limits the ability to generalize the conclusions to a broader biological context. To strengthen these findings, future studies should expand the variety of KRAS^wt^ models and consider using isogenic models, where the only variable difference is the presence of the KRAS mutation.

Absence of Isogenic Models: While comparisons were made between cells with and without KRAS mutations, the study did not include isogenic models in which only the KRAS mutation is altered without affecting other genetic factors. This means that some differences in treatment response could be due to other inherent genetic variations in the cell lines used. Implementing isogenic models in future research would allow for a more precise evaluation of the specific impact of KRAS on therapy resistance.

Variability in Reported Therapeutic Responses in literature: There are discrepancies between our findings and previous studies on the combination of KRAS inhibitors with chemotherapy. While our data suggests synergy between these compounds, previous studies have reported contradictory results, particularly in organoid models or clinical trials with MEK inhibitors. These differences could be attributed to the variability in experimental models used or the specific characteristics of the inhibitors tested. Additional studies directly comparing the effects of different inhibitors in various models will be crucial to clarify these discrepancies.

Lack of In Vivo Validation: Although reference is made to previous studies using animal models, this study does not present in vivo experiments that evaluate the combined efficacy of KRAS inhibitors and chemotherapy in a physiological setting. Validating these findings in animal models would be essential to determine the clinical relevance of the proposed strategy, as well as to explore potential adverse effects or emerging resistance mechanisms in a more complex biological context.

Incompletely Elucidated Molecular Mechanisms: Although a key role of p21 in the induction of senescence-like has been identified, some findings, such as the simultaneous activation of Ki67 and CDK2, suggest the presence of non-conventional mechanisms in cell cycle regulation. Additionally, the relationship between KRAS inhibition, autophagy, and senescence-like has not been fully explored. Future research should delve deeper into these processes to more precisely determine the mechanisms underlying tumor cell responses to treatment.

## Materials and Methods

### Cell lines and cell culture

MiaPaCa-2, Panc-1 and BxPC-3 cells were obtained from the American Type Culture Collection (ATCC, Manassas, VA, USA) and cultured in Dulbecco’s modified Eagle’s medium (DMEM, Thermo Fisher Scientific, 61965-026) containing 10% fetal bovine serum (Hyclone, SV30180.03) in an incubator with 5% CO2 at 37°C, and were authenticated by ATCC as a custom service. Three PDC carrying mutated KRAS (PDAC001T (G12D), PDAC024T (G12D), PDAC054T (G12C) and PDAC022T (G12R)) and one wildtype (PDAC064T) were obtained from patients included in PaCaOmics clinical trial (number 2011-A01439-32). Consent forms of informed patients were collected and registered in a central database. The studies were conducted in accordance with the Declaration of Helsinki.

### Protein extraction and Western blot analysis

To perform protein expression analysis, the cells were detached and homogenized in RIPA buffer. The proteins were separated by SDS-PAGE (29:1 acrylamide: bis-acrylamide, Euromedex Laboratories) in 10% to 12% running gel and 4% stacking gel, in an electrophoresis cell. Proteins were electro-transferred to a nitrocellulose membrane (Immobilon-P, EMD Millipore Corporation) at 250 mA for 2 hours. To identify proteins, the membranes were blocked for 1 hour at room temperature with 5% powdered milk in PBS containing 0.1% Tween 20. Next, they were incubated overnight at 4°C with the rabbit polyclonal antibodies anti-CDK2 (1:1,000; Cell Signaling Technology), anti-CDK4 (1:2,000; Cell Signaling Technology), anti-CDK6 (1:1,000; Cell Signaling Technology), anti-Ki67 (1:1,000; Cell Signaling Technology), anti-p21 (1:1,000; Cell Signaling Technology), anti-p27 (1:1,000; Cell Signaling Technology), anti-ZEB1 (1:1,000; Cell Signaling Technology), anti-Vimentin (1:1,000; Cell Signaling Technology), anti-ERK (1:1,000; Cell Signaling Technology), anti-pERK (1:1,000; Cell Signaling Technology), anti-AKT(1:1,000; Cell Signaling Technology) anti-pAKT S473 (1:1,000; Cell Signaling Technology), anti-pPDK1 S241 (1:1,000; Cell Signaling Technology), anti-pAKT S308 (1:1,000; Cell Signaling Technology), anti-p-PTEN S380 (1:1,000; Cell Signaling Technology), For the immunoreaction, the membranes were incubated with horseradish peroxidase (HRP)-conjugated goat anti-rabbit IgG (1:3,000 dilution, Suther Biotech) or HRP-conjugated rabbit anti-goat IgG (1:3,000 dilution, Suther Biotech). The outcome was visualized using the Chemiluminescent HRP substrates (Millipore Corporation) for chemiluminescence development. To normalize the results, rabbit poly-clonal anti-GAPDH (1:6,000 dilution, Cell Signaling) was used on the same membranes and revealed with HRP-conjugated goat anti-mouse IgG (1:3,000 dilution, Suther Biotech). The membranes were scanned using a PXi multi-application imager (Sygene). The estimation of bands was performed using a prestained protein ladder (SeeBlue Plus2, Thermo Fisher Scientific) as a molecular weight marker. The intensity of each protein band was quantitated using the Fiji (57) software, and the results were expressed as the optical density of each protein/optical density of GADPH.

### Therapy-Induced Senescence Reversion

Five hundred thousand cells per well were seeded in 24-well plates in DMEM or SFDM. Twenty-four hours later, the media were supplemented with increasing concentrations of gemcitabine and incubated for 72 hours. Cellular senescence-like was determinate by β-galactosidase activity with a β-Galactosidase Staining Kit (Cell Signaling). In brief, the media from the cells was removed, washed and fixed for 15 minutes at room temperature. Then, the plates were rinsed again and incubated with β-galactosidase staining solution at 37°C overnight in a dry incubator. To revert this effect, the KRAS G12D inhibitor (MRTX1133) and KRAS G12C (AMG-510) were incubated in increasing concentrations for 3 hours. Then, plates were washed and counterstained with SYBR green (Thermo Fisher Scientific). Images were captured using an optical microscope (Zeiss Axio Imager 2Eclipse 90i; Nikon) with an attached digital camera (DXM1200C; Nikon). The positive and negative cells were counted in three different field using Fiji (57) software applying segmentation by Weka algorithm. The results were represented as percentage of positive cells in relation to the total of cells.

### β-galactosidase expression by luminescence assay

The levels of β-galactosidase were measured using β-galactosidase luminescence assay (Promega). The experiments were performed on a 96-well plate containing DMEM or SFDM. After 24 hours the cells were treated for 72 hours with gemcitabine alone 0.1 µM and 1 µM or in combinatorial scheme of gemcitabine (form 0.01 μM to 10 µM) and the AKT (Ipatasertib) and ERK (SCH772984) inhibitors. To each well 100 µl of Beta-Glo reagent was added and luminescence was measured 1 hour latter using the plate reader Tristar LB941 (Berthold Technologies, Bad Wildbad, Germany). Each experiment was repeated at least three times. Values were normalized and expressed as the fold change of the control (vehicle).

### Immunofluorescence and nuclear area estimation

An amount of 50,000 MiaPaCa-2, Panc-1 and BxPC-3 cells were seeded in twenty-four well plates on coverslips and treated with gemcitabine at 0.1 and 1 µM for 72 hours. After fixation with methanol, the cells were incubated with an anti-rabbit anti-p21 (1:1,000; Cell Signaling Technology) or an anti-rabbit anti-pERK (1:200 Cell Signaling Technology) primary antibody. After washing out the first antibody, cells were incubated with Alexa Fluor 488-labeled anti-rabbit (1:500) secondary antibody (Invitrogen, Barcelona, Spain) and DAPI (4′,6-diamidino-2-phenylindole, Thermo Fisher Scientific, Valencia, Spain) was used to stain the nucleus. Image acquisition was carried out by using a confocal microscope LSM 880 (×63 lens) controlled by Zeiss Zen Black (Zeiss, Oberkochen, Germany). Parallelly, the image from DAPI staining was used to quantify the nuclear area, applying a nuclear mask generate from Fiji software. The estimated area was expressed as the logarithm in base 10.

### Cytotoxicity essays

Three thousand cells per well were seeded in 96-well plates in DMEM or SFDM. Twenty-four hours later, the media were supplemented with increasing concentrations of gemcitabine (from 0.01 μM to 10 µM), MRTX1133 (1 µM), and/or AMG-510 (1 µM) and incubated for 72 hours. It is important to highlight that the effect of the KRAS inhibitors in the absence of gemcitabine is represented at the 0 µM gemcitabine point, allowing us to assess its baseline impact. Cell viability was measured with PrestoBlue (Thermo Fisher Scientific). Fluorescence was measured using the plate reader Tristar LB941 (Berthold Technologies, Bad Wildbad, Germany). Each experiment was repeated at least three times. Values were normalized and expressed as the percentage of the control (vehicle), which represents 100% of normalized fluorescence. Additionally, the cytotoxic activity of the KRAS inhibitors alone was assessed during 72 hours in a range of concentrations between 0.01 μM to 100 μM.

### Time-lapse experiment

Time-lapse analysis was performed by seeding 3,000 cells per well in 96-well plates. After 24 hours, cells were treated with 0.1 μM gemcitabine, 1 μM AMG-510, 1 μM MRTX1133, or a combination of these treatments. A DMSO-treated control was included in parallel. Cell proliferation was monitored using the Incucyte S3 Live-Cell Analysis System (Sartorius) within an incubator maintained at 37 °C in a humidified 5% CO₂ atmosphere. The experiment was conducted over 120 hours (5 days), with images captured every 6 hours. Results are reported as the estimated number of cells per mm².

### Survival analysis and single sample enrichment of bulk RNAseq data

For TCGA-PAAD (n=145) and Puleo (n=288) (23) cohorts the p21 groups were stablished using the maximally selected rank statistical determined by the “surv_cutpoint” function from survminer R package. The top expressed 19% and 20% were identified as the “p21 high group” form TCGA-PAAD and Puleo cohorts, respectively. Kaplan-Meier analysis and Cox proportional-hazards regression model were performed with the Survival R package. Lastly, single sample enrichment analysis of EpiSen signature (24) was performed with the GSVA method using the GSVA R package.

### Single-nucleus data processing and analysis

The single-nucleus data was previously published by Hwang and colleagues (25). The raw counts were processed using Seurat R package. For the analysis of this study only the malignant cells were used. Briefly, the data was normalized, and the 2000 more variable genes were identified with FindVariableFeatures function. After that, the data was scaled, centered, and principal component analysis (PCA) was run with the RunPCA function. The visualization was performed using Uniform Manifold Approximation and Projection (UMAP). EpiSen signature score for each cell was performed with AddModuleScore function.

### Statistical analysis

Statistical analysis comparing two groups was conducted using the unpaired two-tailed Welch’s t-test. For time-lapse experiments were analyzed with 2-ways ANOVA with repetitive measures and the time points were evaluated applying Tukey’s multiple comparisons test. Only the significance of the last time points is reported. The area under the curve (AUC) was computed with the trapezoidal rule using 0 as baseline in Prism 9.5 (Graphpad, California, USA) for the cell viability assays. The results were expressed as the mean ± SD of at least three independent experiments. A *P*<0.05 was regarded as statistically significant.

## Data availability

TCGA-PAAD RNA expression data were downloaded with TCGAbiolinks R package. Puleo cohort data is available in ArrayExpress under the accession number E-MTAB-6134. Processed single-nucleus RNA-seq data is available on the Single Cell Portal under the accession numbers SCP1089 and SCP1096.

## Disclosure of Potential Conflicts of Interest

No potential conflicts of interest were disclosed. The funders had no role in the design of the study; in the collection, analyses, or interpretation of data, in the writing of the manuscript, or in the decision to publish the results.

## Authors’ Contributions

**A. Meilerman**: Resources, data curation, validation, investigation, methodology, writing-review and editing; **P. Santofimia-Castaño**: Conceptualization, formal analysis, validation, investigation; **M. Estaras**: Conceptualization, formal analysis, validation, investigation; **D. Grasso**: Conceptualization, formal analysis, investigation; **E. Chuluyan**: Conceptualization, supervision, investigation; **G. Lomberk**: Conceptualization, formal analysis, validation, investigation, writing-original draft; **R Urrutia**: Conceptualization, formal analysis, validation, investigation, writing-original draft; **N. Dusetti**: Conceptualization, formal analysis, supervision, writing-original draft, writing-review and editing; **N. Fraunhoffer**: Conceptualization, formal analysis, supervision, writing-original draft, writing-review and editing; **J. Iovanna**: Conceptualization, data curation, formal analysis, supervision, funding acquisition, writing-original draft, project administration, writing-review and editing.

## Acknowledgments

This work was supported by INCa (grants number 2018-078 and 2018-079), Canceropole PACA and INSERM.

**Legend of Supplementary Figure 1**

**Increase of SA-β-gal levels after chemotherapeutic treatment in primary PDAC-derived cells.** A. Induction of SA-β-gal after the treatment with gemcitabine (72 hours) 0.1 µM and 1 μM. B. Percentage of SA-β-gal cell after the treatment with gemcitabine at 0.1 and 1 µM for 72 hours. Scale bars, 100 μm. \**P*<0.05, \*\**P*<0.01, \*\*\**P*<0.001. (n = 3).

**Legend of Supplementary Figure 2**

**Cytotoxic profile of KRAS inhibitors MRTX1133 and AMG-510.** Three commercial cell lines MiaPaCa-2, Panc-1, and BxPC-3 and four patient-derived primary cell cultures were treated with increased concentrations of MRTX1133 and AMG-510 (from 0.01 μM to 100 μM) for 72 hours. Cell viability was measured with PrestoBlue. Each experiment was repeated at least three times. Values were normalized and expressed as the percentage of the control (vehicle).

## Notes

### Competing Interest Statement

The authors have declared no competing interest.

