## Supplementary figures and images for "KRAS Inhibition Reverses Chemotherapy Resistance Promoted by Therapy-Induced Senescence-like in Pancreatic Ductal Adenocarcinoma"

### Supplemental Figure 1

#### Slide 1
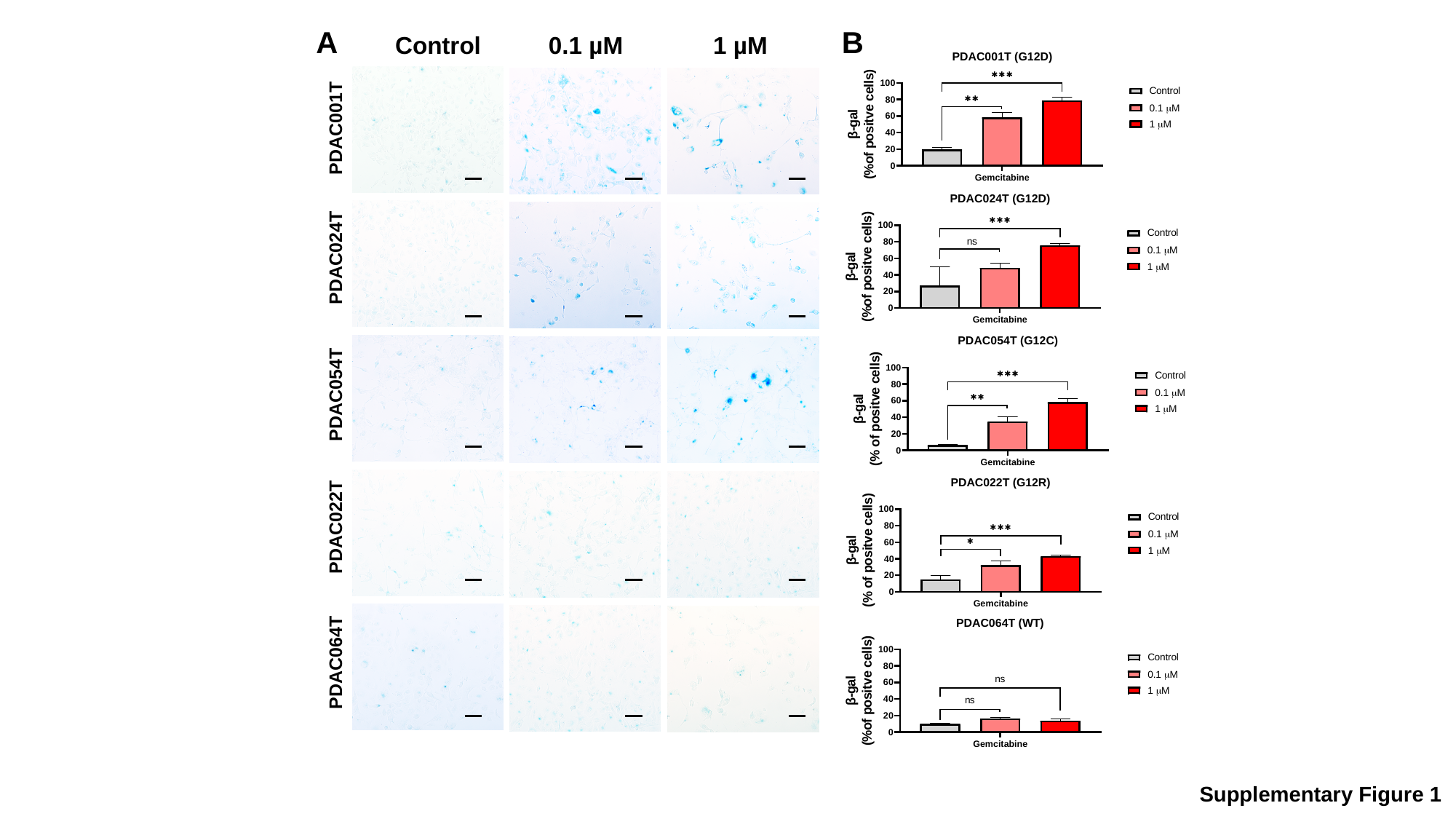

A
B
Control
0.1 µM
1 µM
PDAC001T
PDAC024T
PDAC054T
PDAC022T
PDAC064T
Supplementary Figure 1

### Supplemental Figure 2

#### Slide 1
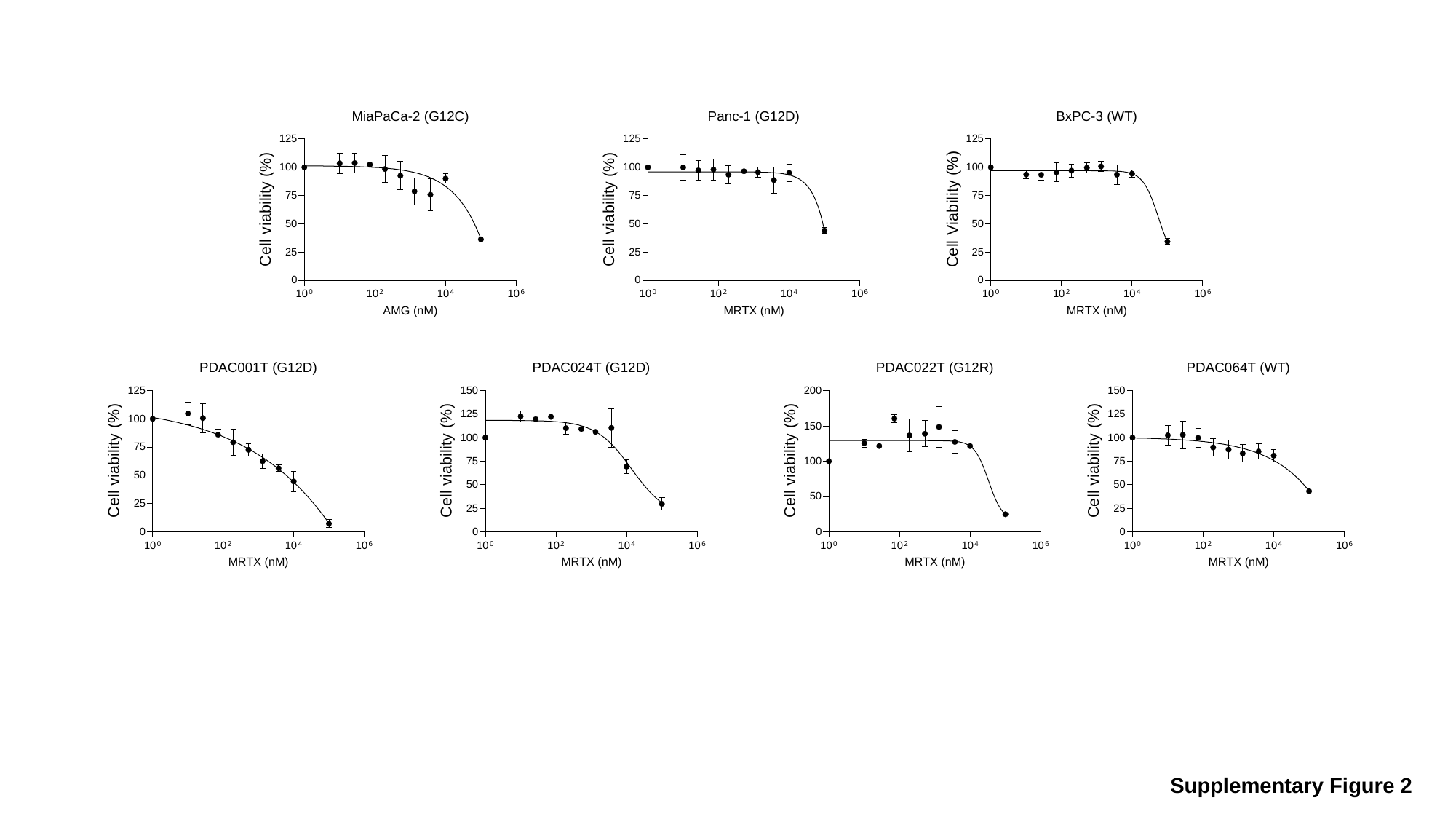

Supplementary Figure 2
